# mOTUpan: a robust Bayesian approach to leverage metagenome assembled genomes for core-genome estimation

**DOI:** 10.1101/2021.06.25.449606

**Authors:** Moritz Buck, Maliheh Mehrshad, Stefan Bertilsson

## Abstract

Recent advances in sequencing and bioinformatics have expanded the tree of life by providing genomes for uncultured environmentally relevant clades, either through metagenome assembled genomes (MAGs) or single-cell assembled genomes (SAGs). While this expanded diversity can provide novel insights about microbial population structure, most tools available for core-genome estimation are overly sensitive to genome completeness. Consequently a major portion of the huge phylogenetic diversity uncovered by environmental genomics approaches remain excluded from such analyses. We present mOTUpan, a novel iterative Bayesian method for computing the core-genome for sets of genomes of highly diverse completeness range. The likelihood for each gene-cluster to belong to core- or accessory-genome is estimated by computing the probability of its presence/absence pattern in the target set of genomes. The core-genome prediction is computationally efficient and can be scaled up to thousands of genomes. It has shown comparable estimates as state-of-the-art tools Roary and PPanGGoLiN for high-quality genomes and outperforms them at lower completeness thresholds. mOTUpan wraps a bootstrapping procedure to estimate the quality of a specific core-genome prediction, as the accuracy of each run will depend on the specific completeness distribution and the number of genomes in the dataset under scrutiny. mOTUpan is implemented in the mOTUlizer software package, and available at github.com/moritzbuck/mOTUlizer, under GPL 3.0 license.

## Introduction

The continuous advancements of high throughput sequencing technologies and bioinformatics tools over the last two decades have fueled large-scale ecogenomics analyses leading up to a new view of the tree of life[1, 2, 3]. This refined view enabled by metagenomics and single-cell genomics reveals that uncultured bacteria and archaea exclusively represented by MAGs (metagenome assembled genome) and SAGs (single-cell amplified genome) account for ca. 75% of the cataloged phylogenetic microbial diversity[2]. Despite their unequivocal potential to reveal diversity, the inherent incompleteness of MAGs and SAGs has so far hindered attempts in the large-scale study of sub-population diversity, core-genome structure, and genome evolution of these phylogenetically diverse species.

All non-redundant genes in genomes from a genome-set are part of its pangenome and can be categorized as either core or accessory[4]. The core-genome is a set of genes common among all genomes of a species and are supposedly responsible for the basic aspects of the cell’s biology and phenotypic traits[5]. The accessory part of the genome is underpinning the sub-species diversity and is defined as genes present in two or more but not all representatives of a species. Accessory genes typically encode for functions that provide cells with adaptive advantages (e.g., supplementary metabolic pathways, enzymatic activities, antibiotic resistance, phage and predation resistance, pathogenicity, etc.)[4, 5, 6], but are often also relics or live selfish genetic elements [7].

A key prerequisite for the comparative analyses of the sub-species diversity and ecological adaptations is to first have a robust estimation of the core-genome that will enable a better assessment of the accessory counterparts. However, core-genome analyses are limited in taxonomic scope[8, 9, 10, 11, 12, 13], largely because of the severe limitations in culturing microbes and obtaining high quality genomes, combined with existing bioinformatics methods being dependent on high quality genomes to scaffold such analyses. Most methods used for core-genome analysis only work with sets of high quality and complete genomes and are very sensitive to missing genes and fragmented genomes[14]. These methods often concentrate on developing novel methods for computation of clusters of orthologous genes (COGs) in the population of interest[14] and use only simple binary presence/absence models for the core-genome estimation (e.g. a COG is core if it is in all of the genomes of the clade). Such methods perform best when used on a moderate number of high-quality genomes generated from cultured microbial isolates. Accordingly, these methods are unable to deal with the rapidly growing database of incomplete and fragmented MAGs and SAGs of the uncultured majority of earth’s microbiome[2]. Due to these methodological limitations, our understanding of the size and structure of microbial core-genomes and pangenome dynamics remain elusive and lag behind our growing appreciation of microbial phylogenetic diversity. The recently released software, PPanGGoLiN, uses synteny networks to compute clusters of co-occurring gene-clusters instead of presence/absence. This method is highly scalable, fast, and robust enough to deal with incomplete genomes[15]. However, this method could be sensitive to fragmentation which is a prominent feature of most incomplete MAGs and SAGs, and is not explicitly tailored to find the core, but rather to find clusters of synthenic genes.

Here we present a novel approach for computing core-genomes relying on a Bayesian estimator of the observed presence/absence patterns of discrete genome-encoded traits (any trait that can be encoded in a genome, e.g. gene-cluster, COG, functional annotations, etc.) in sets of incomplete MAGs/SAGs and complete genomes. We wrote a software tool, mOTUpan, that can estimate if any genome-encoded trait is more likely to be present in all genomes of a genome-set or only in a subset. mOTUpan can compute the core-genome partitioning for genome-sets of a wide range of qualities, and is computationally efficient, agnostic to the genome-encoded traits used, and very robust to incompleteness.

## Methods

### Bayesian approach for core-genome estimation

mOTUpan can use any set of genomes that is suspected to share a certain number of genome-encoded traits. We typically use clusters where all genomes are within compact clusters defined by a 95% average nucleotide identity (ANI) threshold. We call such clusters mOTUs (metagenomic Operational Taxonomic Units), that can be seen as an operational definition of species. However, genomes clustered at any other taxonomic level, or any other way one can imagine (by niche, predator, host, etc.), are valid too. We will use the term genome as a shorthand for any set of nucleotide sequences originating from the same organism. This could be draft genomes, complete genomes, MAGs, or SAGs. Each genome is first described as a set of genome-encoded traits. Here we will use gene-clusters, but it should be mentioned that mOTUpan is agnostic to the specific form of such traits, one could use genes, COGs, functional annotations, or any other discrete trait that is encoded by a genome. mOTUpan then uses an iterative Bayesian approach to classify each trait of the genome in a genome-cluster as a core- or accessory-trait based on a likelihood ratio. For each of the two hypotheses (core- or accessory-trait) a probability is computed using a genome completeness prior inferred for each genome (genome completeness can be calculated using CheckM[16], or a fixed value used). The most likely trait category (core or accessory) is then picked as class for that trait. Using this new classification, we compute posterior completeness estimates, which can be used as a prior for a second iteration and then repeat this entire process until convergence.

### Probability models

To compute the probability of a distribution of a specific trait in the genome-set mOTU under the assumption that it is in the core, we multiply the propability *p*_trait∈g|core_ of any genome *g* (*g* is treated as a set of traits) that has that gene-cluster, with the inverse probability 1 *−p*_trait∈g|core_ for the genomes that do not have that trait. Where the probability *p*_trait∈g|core_ is actually directly the prior completeness estimate *c_g_* of *g*, e.g. equations 1 and 2:

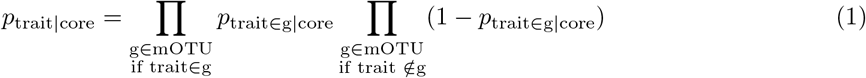

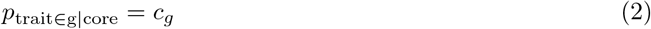

For the probability under the assumption that it is in the accessory fraction of the genome, we will have to make some further assumptions with regards to the structure of the pangenome. We have assumed that the traits in the pangenome that are not in the core, are independent, and each trait has a frequency 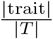 where |trait| is the number of genomes in mOTU that have that trait, and *|T |* the total size of the traits-pool, e.g. Σ_all traits_ |trait|. To “fill” the accessory fraction of a genome, we draw “*|g| − c_g_|*core_mOTU_|”-times, which is the number of spots in the accessory part of the genome assuming a genome with |*g*| traits, core size |core_mOTU_| and completeness *c_g_*. Resulting in equations 3 and 4:

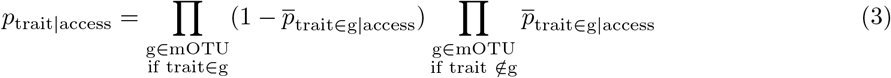

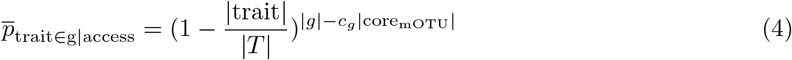

For practical reasons, these computations are all done in log-space, resulting in a log-likelihood ratio:

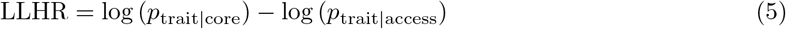

If the LLHR of equation 5 is positive, the trait is considered core, if negative, it is considered accessory. Using this classification, we recompute an updated completeness estimate for each genome:

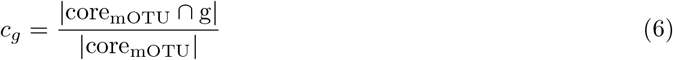

where core_mOTU_ is the set of all traits classified as core.

After this step, we rerun the likelihood computation. This is repeated until convergence, to obtain a final set of core-traits and accessory-traits, and posterior completeness estimates.

### Bootstrapped false-discovery rate and sensitivity

In addition to the likelihood ratio between the two probabilities, a bootstrapping approach has been integrated in mOTUpan to estimate the false-discovery rate and sensitivity of a specific partitioning. Synthetic genomes are built by drawing gene-clusters from the original genome-set according to the partitioning (e.g. every synthetic genome has all of the core genes-clusters, and the accessory gene-clusters are drawn randomly from the other gene-clusters according to their frequency). These synthetic genomes are then rarefied according to the genome-set’s posterior completeness estimate. This synthetic set of genomes are then run through mOTUpan again and these results are used to estimate the false-positive rate and sensitivity. Multiple synthetic data-sets can be analyzed to obtain a better estimate.

### Benchmarking mOTUpan for core-genome estimation

To benchmark the core-genomes computed by mOTUpan against other commonly used core-genome analysis tools, we calculated the core-genomes for 301 species containing a total of 11570 genomes (species were rarefied to 50 genomes to make the runs tractable with Roary) from the genome taxonomy database (GTDB)[3] and 258 mOTUs containing 8955 genomes in total from the StratFreshDB[17]. The MAGs were reclustered with mOTUlize[18] with less stringent parameters (“`--MAG-completeness 30--MAG-contamination 10`”) to have more low quality mOTUs and compare the performance of mOTUpan to Roary[14] and PPanGGOLiN[15]. Genome statistics, accession numbers and taxonomy are available in the Supplementary Table S1. This step aims to highlight and compare the performance of mOTUpan with Roary and PPanGGOLiN with regards to the ability to handle incomplete and fragmented genomes.

For more detailed benchmarking of mOTUpan performance, we selected a dataset of genomes affiliated with the *Prochlorococcus_A* genus from the GTDB. All genomes classified as *Prochlorococcus_A* according to GTDBtk[19] found in RefSeq as well as GORG[20], were clustered into mOTUs (using mOTUlize[18] with standard parameters), the mOTU with the largest number of genomes was used (Supplementary Table S2 for genome statistics and accession numbers). This *Prochlorococcus* mOTU consists of 388 genomes whereof 3 are closed genomes and 16 genomes are estimated to be more than 95% complete according to CheckM[16] results. Genomes assigned to this mOTU range in completeness from 8.59% to 99.52% (median=69.05%) (Supplementary Table S2). mOTUpan’s performance for core-genome estimates for this *Prochlorococcus* mOTU was benchmarked against PPanGGOLiN using the gene-clusters generated by it (PPanGGOLiN uses mmseqs[21] internally for gene-clustering).

## Results and Discussion

### Overview of the mOTUpan’s Bayesian approach

The Bayesian approach adopted in this tool allows us to leverage the genomic diversity uncovered by incomplete and fragmented MAGs and SAGs for exploring the core-genome and pangenome structure of bacterial and archaeal species (or any other set of genomic traits). Most available tools such as Roary rely on a hard presence/absence threshold for defining the core-genome. This limitation renders such tools largely unusable when dealing with incomplete and fragmented MAGs and SAGs. Comparing the performance of Roary and mOTUpan for core-genome estimation with the gene-clusters computed by Roary is equivalent to comparing mOTUpan to a hard threshold approach.

The network nature of PPanGGOLiN makes it relatively robust to deal with some degree of incompleteness, however as it is looking for patterns of synteny to determine the persistent fraction of the genomes, too much fragmentation (that is common in MAGs and SAGs) could cause problems in calculations of the persistent fraction of the genomes. In the case of species represented by MAGs and SAGs, the genes that are classified by PPanGGOLiN as “persistent” are very likely to be a part of the core, but the approach will likely overlook some core genes. The genes classified as “shell” will thus contain part of the core-genome as well as highly prevalent genes often organized as operons. mOTUpan on the other hand, bypasses both incompleteness and fragmentation limitations and offers a robust estimation of the core-genome and pangenome for sets of incomplete and fragmented MAGs and SAGs. mOTUpan also calculates bootstrapped false-discovery rate and sensitivity for the core-genome/pan-genome partitioning.

There are widespread and valid concerns that MAGs are contaminated by contigs that might not be a genuine part of their genome, as binning tools may mistakenly cluster them together with the rest of the MAG. MAGs are usually screened for putative contamination with tools such as CheckM that relies on a limited dataset of high-quality genomes to compute a set of markers. mOTUpan can however address this known problem in a different way, as genes annotated as core have a very low likelihood of being contaminants and can thus be used for prediction of genome quality. Thus, mOTUpan allows users to refine the completeness values estimated by CheckM. Additionally, for other genome-sets, viruses or plasmids, mOTUpan can still obtain a completeness estimate.

### Benchmarking mOTUpan against Roary and PPanGGOLiN along the completeness scale

To benchmark the performance of mOTUpan against Roary, we used the gene-clusters generated by Roary. Comparing the performance along the completeness scale shows that Roary is highly sensitive to genome completeness, as Roary’s core-genome estimate drop away considerably from that of mOTUpan when completeness decreases (Fig. 1A-B). Some of these limitations can be bypassed by manually adjusting thresholds in Roary, but while this can be done at a small scale, it is not tractable for the larger scales where mOTUpan can still function (as is stated on its web-page^1^ “Roary is not intended for meta-genomics or for comparing extremely diverse sets of genomes”’).

**Figure 1.**
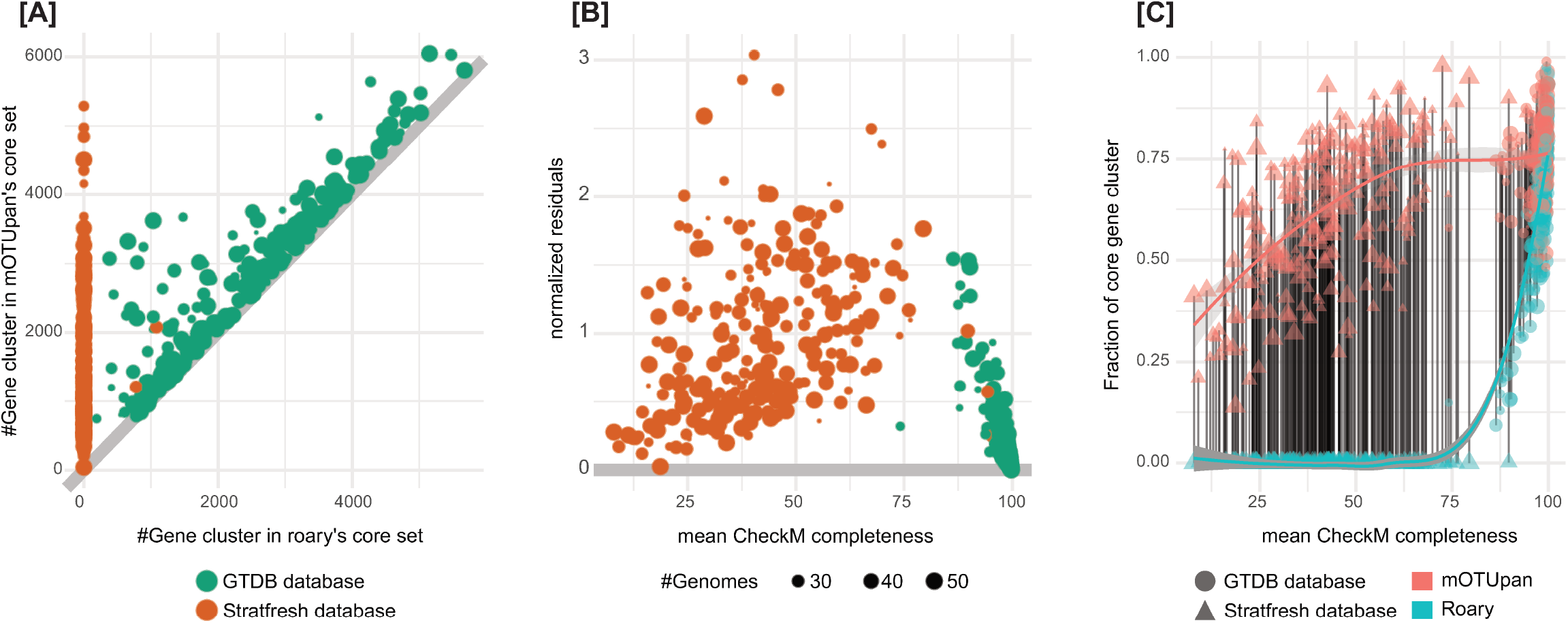
Benchmarking the performance of mOTUpan against Roary along the completeness scale. 301 species containing 11570 genomes from the genome taxonomy database (GTDB) and 258 mOTUs containing 8955 genomes in total from the StratfreshDB are used for this comparison. Gene-clusters used are the ones computed by Roary. A) Predicted core sizes. B) normalized residues, fold change between core size predicted by mOTUpan and roary, if the number is larger than one, mOTUpan’s prediction is larger. C) Fraction of genome made of gene-clusters in the core.

Running mOTUpan using the COGs generated by PPanGGOLiN (which internally uses the mmseq2[15] clustering tool), we obtain similar core-genome estimates for the GTDB data-set (the more complete genome-sets) (Fig.2A). Looking more specifically at the deviation from the first bisector along the completeness scale (Fig.2B), we can see that in general PPanGGOLiN’s core-genome estimates are larger than those obtained with mOTUpan for the more complete genome-sets. This tendency changes drastically once the average completeness drops below 70% where the mOTUpan estimates become larger. This increase could be due to an inflation of predicted core gene-clusters for the more incomplete genome-sets. We accounted for this possibility by inspecting the fraction of the genome classified as core (Fig.2C). While this estimate is expected to be independent of completeness, we can see that output from both PPanGGOLiN and mOTUpan drop away from the expected value with lower completeness, but the output from PPanGGOLiN drops faster, demonstrating mOTUpan’s robustness to incomplete and noisy genomes.

**Figure 2.**
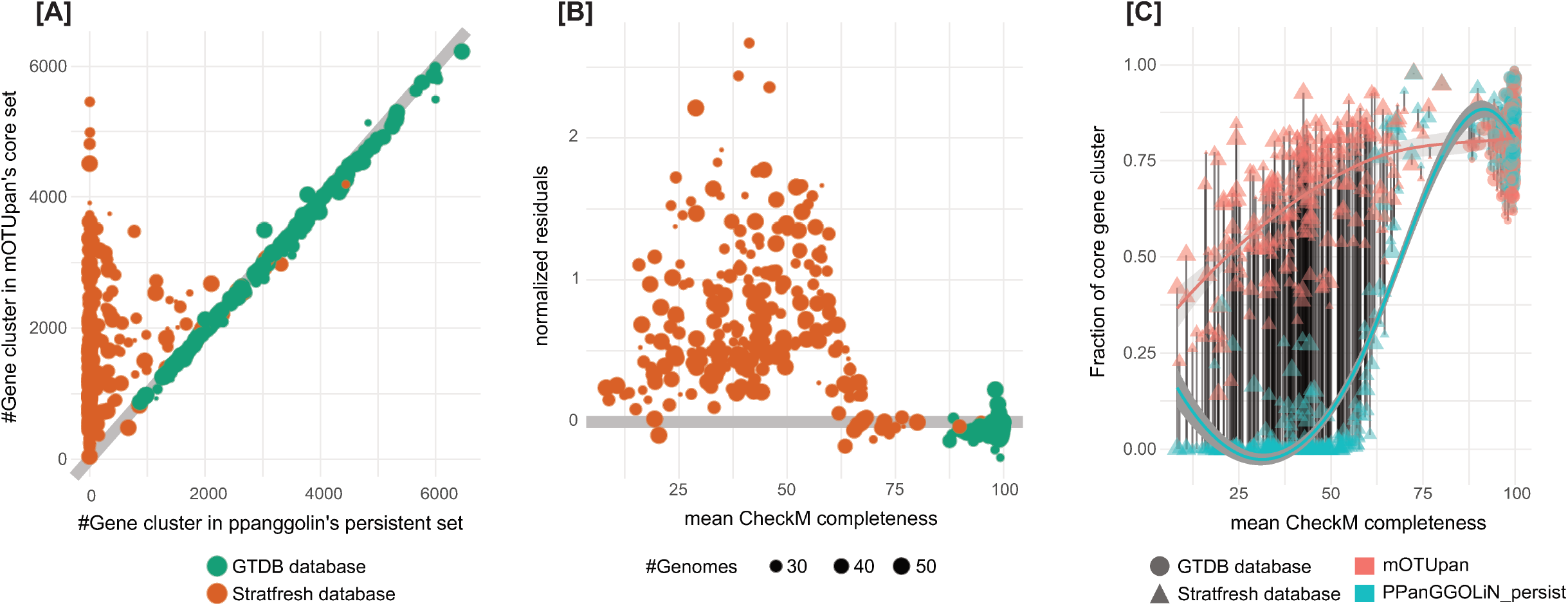
Benchmarking the performance of mOTUpan against PPanGGOLiN along the completeness scale. 301 mOTUs containing 11570 genomes from the genome taxonomy database (GTDB) and 258 mOTUs containing 8955 genomes in total from the StratfreshDB were used for this comparison. Gene-clusters used are the ones computed by PpanGGOLiN (based on mmseqs2). A) Predicted core sizes. B) normalized residues, fold change between core size predicted by mOTUpan and Roary, if the number is larger than one, mOTUpan’s prediction is larger. C) Fraction of genome made of gene-clusters in the core.

### Benchmarking mOTUpan against PPanGGOLiN for a *Prochlorococcus_A* genome-set

For a more detailed benchmarking of mOTUpan against PPanGGOLiN, we used a set of 388 genomes from the *Prochlorococcus_A* genus, ranging in completeness from 8.59% to 99.52% (median=69.05%) according to CheckM (Supplementary Table S2). For this analysis we used the gene-clusters generated by PPanGGOLiN.

PPanGGOLiN splits the set of gene-clusters by default into three subsets: persistent, shell and cloud. For very complete genomes, the persistent set of gene-clusters is close to the core-genome, but for more noisy genomes, such as those included in this *Prochlorococcus_A* genome-set, the approach is not capturing the entire core-genome (Fig.3). It is notable that gene-clusters identified as “persistent” (316 gene-clusters) very likelily belong to the core-genome while the “shell”-set of genes will normally correspond to frequently co-occurring genes. PPanGGOLiN estimates a total of 1537 gene-clusters to be a part of the “shell” category for the *Prochlorococcus_A* gene-set. For the same gene-set, mOTUpan estimates 1637 gene-clusters to be part of the core-genome. The core estimate of mOTUpan seems to be close to the sum of “persistent” and “shell” (1853 gene-clusters). The three closed genomes have 1883 gene-clusters, making the “persistent+shell” estimate probably an overestimate of the core-genome. The “shell” set of gene-clusters is picking up genes that are probably not all from the core but rather frequently occurring accessory operons. This is shown in the heatmap in Fig.4. Conversely, it also shows the robustness of mOTUpan to estimate the true core-genome from more noisy mOTUs.

**Figure 3.**
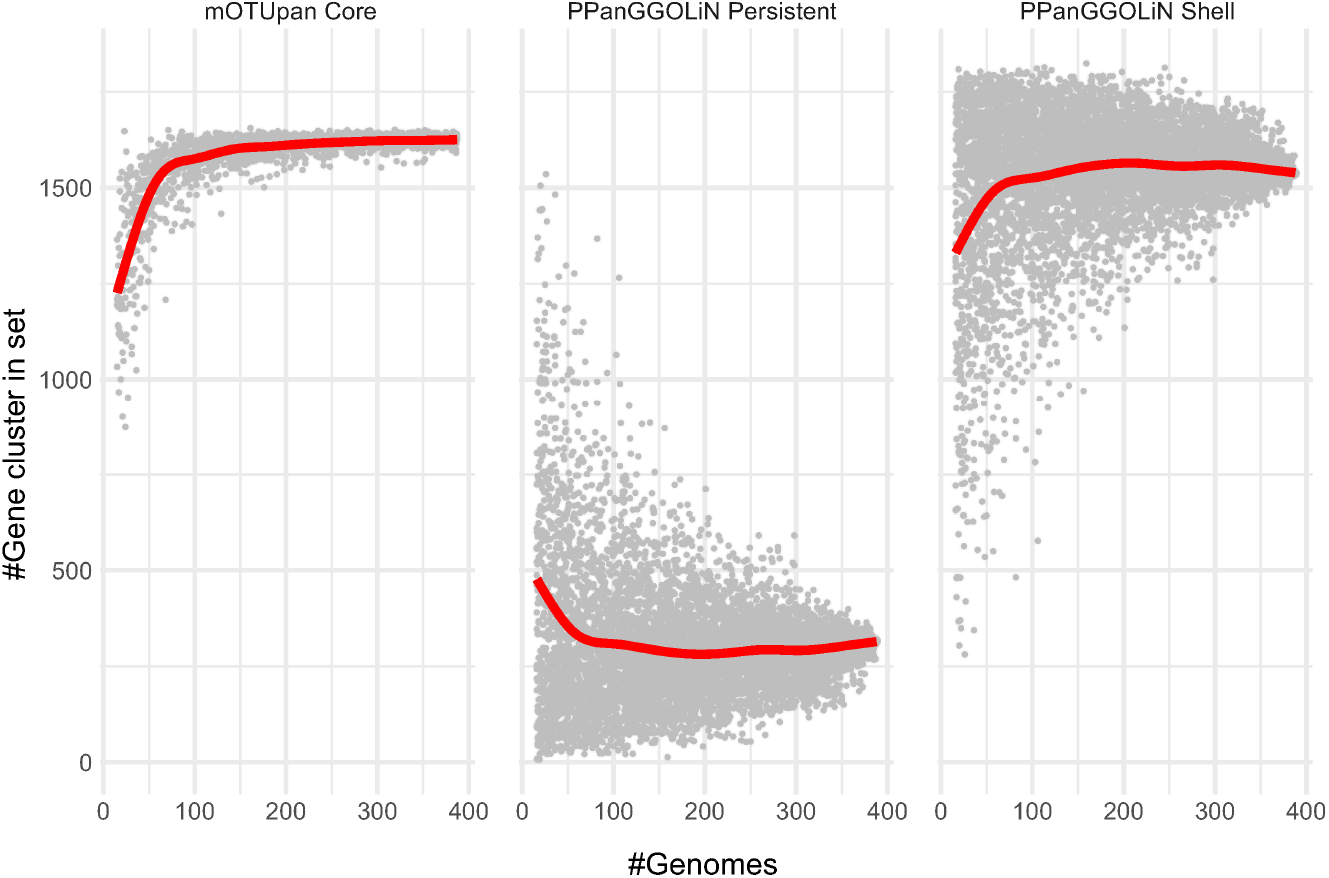
Rarefaction analysis of mOTUpan’s and PPanGGOLin’s core-genome prediction on the *Prochlorococcus_A* mOTU. The same analysis was performed on random subsets of the available 388 genomes.

**Figure 4.**
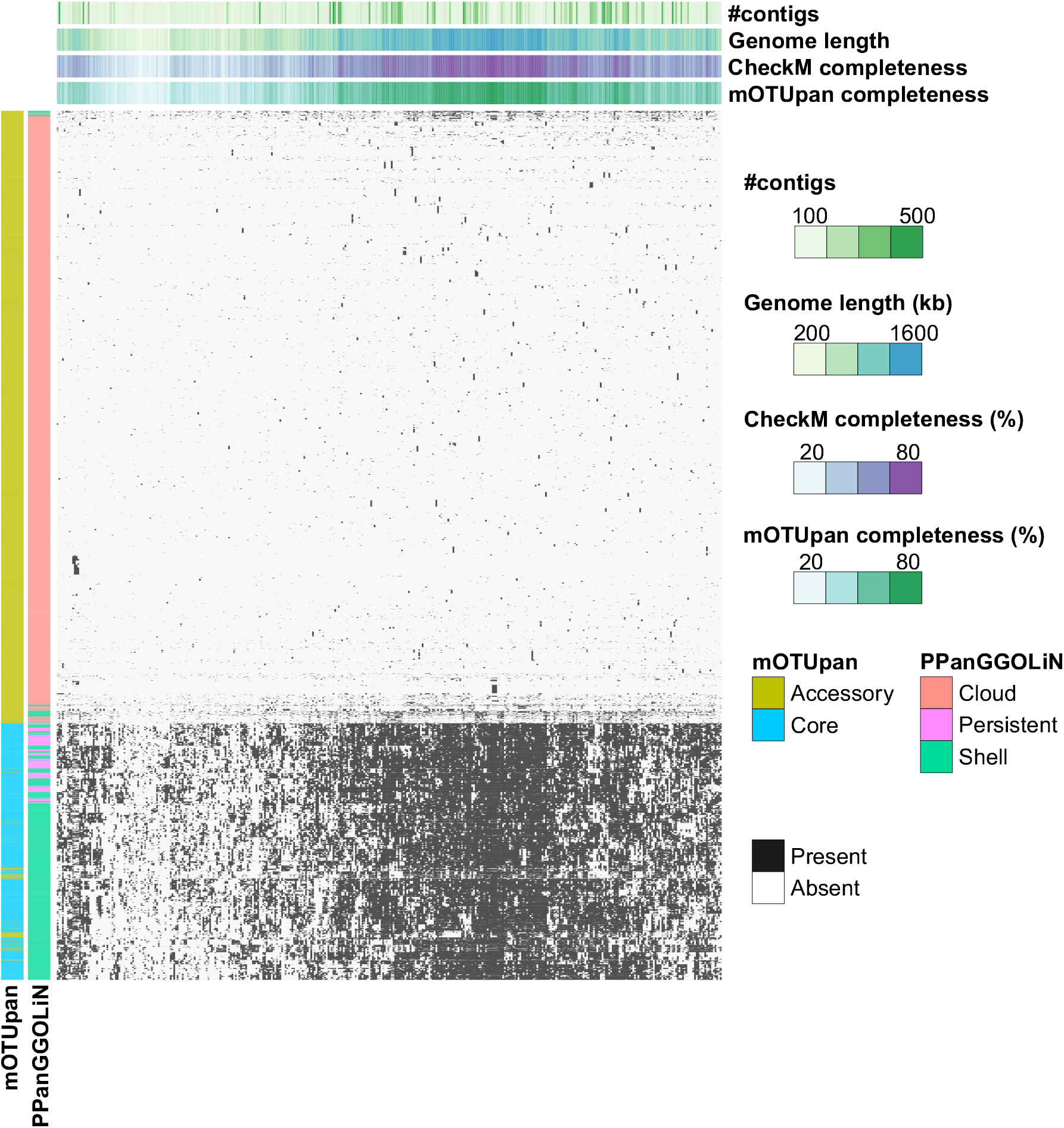
Distribution of 5985 generated gene-clusters in 388 genomes of a *Prochlorococcus_A* mOTU. Columns are genomes, and rows represent gene-clusters. Gene-clusters are assigned to different partitions using mOTUpan and PPanGGOLiN estimations (colored columns on the left). The gene-clusters, that mOTUpan called as accessory and PPanGGOLiN called as shell, seem to belong to blocks of gene-clusters absent in sets of highly complete genomes, hinting at very prevalent operons of accessory genes. Conversely gene-clusters in mOTUpan’s accessory and PPanGGoLiN’s shell, seem to be very prevalent gene-clusters that have only a diffuse pattern hinting at single mobile genes, for example.

Calculations of the core-genome using mOTUpan with the 3 closed genomes and 16 genomes with completeness higher than 95% of the *Prochlorococcus_A* cluster estimates 1644 gene-clusters in the core (1714 “persistent” gene-clusters with PPanGGOLiN). This is probably an upper-bound to the size of the core of this *Prochlorococcus_A* mOTU, as additional micro-diversity and noise would only remove genes from this, making the 1637 gene-clusters predicted to make up the core in mOTUpan for the full set a better estimate than either PPanGGOLiN’s “shell”-set (316 clusters) or “persistent+shell”-set (1853 clusters).

This generally shows that mOTUpan can predict a core-genome very similar to other state-of-the-art tools, while at the same time being more robust over broader ranges of genome completeness in comparison to those tools.

mOTUpan can be used in a number of ways. It can obviously be used to study pan-genome structure at large scale and with noisier data. This comes with some caveats, i.e. the method is highly dependent on the gene-clustering method used and it is very hard to evaluate the correctness of these at a larger scale. Additionally, mOTUpan can only classify genes that actually are in the genomes that are analyzed. Accordingly, genes that are hard to assemble or bin (due to different k-mer or abundance profiles) will be overlooked, leading to an inevitable underestimate of the accessory genomes. Nevertheless, it is the only tool available that can do this type of analysis, and should hence be an invaluable resource for biodiversity exploration and comparative genomics. While PPanGGoLiN is performing very well with noisy data, the specific purpose and scope of this tool is different. PPanGGoLiN can be leveraged if one needs to select and identify core genes to e.g. make a core phylogeny, but mOTUpan is a better choice for estimating and exploring the core- and/or accessory genome structure. Another important use envisioned for mOTUpan is to strengthen functional predictions for metagenomic projects. Rather than relying on single MAGs where the presence of specific genes can be questioned, mOTUpan can robustly quantify this presence as long as highly similar MAGs are available (which is often the case in medium-to-large scale metagenomic project). Notably, it can be used with a variety of genome-encoded traits, and the currently available version has parsers available for: Roary, PPanGGoLiN, eggNOGmapper[22], and mmseqs2[21], with possibly more to be included later.

Ultimately, mOTUpan introduces and enables a new type of analysis within the field of microbial genomics, i.e. the usage of presence-absence of genome-encoded traits combined with some Bayesian computation to predict gene-content in a genome-set. This approach can be expanded into a number of different directions. We can for example move from presence-absence to gene-count, or use this approach for gene-linkage assessment to estimate if some traits co-occur more often than by chance.

## Acknowledgments

Bioinformatics analyses were carried out utilizing the Uppsala Multidisciplinary Center for Advanced Computational Science (UPPMAX) at Uppsala University under projects SNIC 2020/5-19 and 2021/5-53. Funding was provided by the Swedish Research Council (grant 2017-04422 and 2018-04685). Also big thanks to Julia Nuy and Matthias Hötzinger for some early testing.

## Data and code availability

The mOTUpan software is written in Python 3 and is freely available under GPL 3.0 license via GitHub in the mOTUlizer package at github.com/moritzbuck/mOTUlizer. A conda recipe and pip package for user friendly installation are also available in the appropriate repository. Scripts used for the analyses in this paper can be found at github.com/moritzbuck/mOTUlizer/tree/master/mOTUlizer/scripts The data used for benchmarking is from the GTDB[3](release 95), available at gtdb.ecogenomic.org (with actual genomes at RefSeq and Genbank); GORG-Tropics[20], available under GenBank at PRJEB33281; and the StratFreshDB[17]

## Footnotes

1 https://sanger-pathogens.github.io/Roary/

## References

[1] Laura A. Hug, Brett J. Baker, Karthik Anantharaman, Christopher T. Brown, Alexander J. Probst, Cindy J. Castelle, Cristina N. Butterfield, Alex W. Hernsdorf, Yuki Amano, Kotaro Ise, Yohey Suzuki, Natasha Dudek, David A. Relman, Kari M. Finstad, Ronald Amundson, Brian C. Thomas, and Jillian F. Banfield. A new view of the tree of life. Nature Microbiology, 1(5):1–6, April 2016. ISSN 2058–5276. doi: 10.1038/nmicrobiol.2016.48.

[2] Stephen Nayfach, Simon Roux, Rekha Seshadri, Daniel Udwary, Neha Varghese, Frederik Schulz, Dongying Wu, David Paez-Espino, I.-Min Chen, Marcel Huntemann, Krishna Palaniappan, Joshua Ladau, Supratim Mukherjee, T. B. K. Reddy, Torben Nielsen, Edward Kirton, José P. Faria, Janaka N. Edirisinghe, Christopher S. Henry, Sean P. Jungbluth, Dylan Chivian, Paramvir Dehal, Elisha M. Wood-Charlson, Adam P. Arkin, Susannah G. Tringe, Axel Visel, Tanja Woyke, Nigel J. Mouncey, Natalia N. Ivanova, Nikos C. Kyrpides, and Emiley A. Eloe-Fadrosh. A genomic catalog of Earth’s microbiomes. Nature Biotechnology, 39(4):499–509, April 2021. ISSN 1546–1696. doi: 10.1038/s41587-020-0718-6.

[3] Donovan H. Parks, Maria Chuvochina, Pierre-Alain Chaumeil, Christian Rinke, Aaron J. Mussig, and Philip Hugenholtz. A complete domain-to-species taxonomy for Bacteria and Archaea. Nature Biotechnology, 38(9):1079–1086, September 2020. ISSN 1546–1696. doi: 10.1038/s41587-020-0501-8.

[4] Michael A. Brockhurst, Ellie Harrison, James P. J. Hall, Thomas Richards, Alan McNally, and Craig MacLean. The Ecology and Evolution of Pangenomes. Current Biology, 29(20):R1094–R1103, October 2019. ISSN 0960–9822. doi: 10.1016/j.cub.2019.08.012.

[5] Duccio Medini Claudio Donati, Hervé Tettelin, Vega Masignani, and Rino Rappuoli. The microbial pan-genome. Current Opinion in Genetics & Development, 15(6):589–594, December 2005. ISSN 0959-437X. doi: 10.1016/j.gde.2005.09.006.

[6] Maria Rosa Domingo-Sananes and James O. McInerney. Mechanisms That Shape Microbial Pangenomes. Trends in Microbiology, 29(6):493–503, June 2021. ISSN 0966-842X, 1878-4380. doi: 10.1016/j.tim.2020.12.004.

[7] Rosario Gil and Amparo Latorre. Factors Behind Junk DNA in Bacteria. Genes, 3(4):634–650, December 2012. doi: 10.3390/genes3040634.

[8] Steven J. Biller, Paul M. Berube, Debbie Lindell, and Sallie W. Chisholm. Prochlorococcus: The structure and function of collective diversity. Nature Reviews Microbiology, 13(1):13–27, January 2015. ISSN 1740–1534. doi: 10.1038/nrmicro3378.

[9] Yongjun Fang Zhaolong Li, Jiucheng Liu, Changlong Shu, Xumin Wang, Xiaowei Zhang, Xiaoguang Yu, Duojun Zhao, Guiming Liu, Songnian Hu, Jie Zhang, Ibrahim Al-Mssallem, and Jun Yu. A pangenomic study of Bacillus thuringiensis. Journal of Genetics and Genomics, 38(12):567–576, December 2011. ISSN 1673–8527. doi: 10.1016/j.jgg.2011.11.001.

[10] Ryan A. Blaustein, Alexander G. McFarland, Sarah Ben Maamar, Alberto Lopez, Sarah Castro-Wallace, and Erica M. Hartmann. Pangenomic Approach To Understanding Microbial Adaptations within a Model Built Environment, the International Space Station, Relative to Human Hosts and Soil. mSystems, 4(1), January 2019. ISSN 2379–5077. doi: 10.1128/mSystems.00281-18.

[11] Tom O. Delmont and A. Murat Eren. Linking pangenomes and metagenomes: The Prochlorococcus metapangenome. PeerJ, 6:e4320, January 2018. ISSN 2167–8359. doi: 10.7717/peerj.4320.

[12] Mario López-Pérez and Francisco Rodriguez-Valera. Pangenome Evolution in the Marine Bacterium Alteromonas. Genome Biology and Evolution, 8(5):1556–1570, April 2016. ISSN 1759–6653. doi: 10.1093/gbe/evw098.

[13] Philippe Deschamps Yvan Zivanovic, David Moreira, Francisco Rodriguez-Valera, and Purificación López-García. Pangenome evidence for extensive interdomain horizontal transfer affecting lineage core and shell genes in uncultured planktonic thaumarchaeota and euryarchaeota. Genome Biology and Evolution, 6(7):1549–1563, June 2014. ISSN 1759–6653. doi: 10.1093/gbe/evu127.

[14] Andrew J. Page, Carla A. Cummins, Martin Hunt, Vanessa K. Wong, Sandra Reuter, Matthew T. G. Holden, Maria Fookes, Daniel Falush, Jacqueline A. Keane, and Julian Parkhill. Roary: Rapid large-scale prokaryote pan genome analysis. Bioinformatics (Oxford, England), 31(22):3691–3693, November 2015. ISSN 1367–4811. doi: 10.1093/bioinformatics/btv421.

[15] Guillaume Gautreau Adelme Bazin, Mathieu Gachet, Rémi Planel, Laura Burlot, Mathieu Dubois, Amandine Perrin, Claudine Médigue, Alexandra Calteau, Stéphane Cruveiller, Catherine Matias, Christophe Ambroise, Eduardo P. C. Rocha, and David Vallenet. PPanGGOLiN: Depicting microbial diversity via a partitioned pangenome graph. PLOS Computational Biology, 16(3):e1007732, March 2020. ISSN 1553–7358. doi: 10.1371/journal.pcbi.1007732.

[16] Donovan H. Parks, Michael Imelfort, Connor T. Skennerton, Philip Hugenholtz, and Gene W. Tyson. CheckM: Assessing the quality of microbial genomes recovered from isolates, single cells, and metagenomes. Genome Research, 25(7):1043–1055, July 2015. ISSN 1549–5469. doi: 10.1101/gr.186072.114.

[17] Moritz Buck Sarahi L. Garcia, Leyden Fernandez, Gaëtan Martin, Gustavo A. Martinez-Rodriguez, Jatta Saarenheimo, Jakob Zopfi, Stefan Bertilsson, and Sari Peura. Comprehensive dataset of shotgun metagenomes from oxygen stratified freshwater lakes and ponds. Scientific Data, 8(1):131, May 2021. ISSN 2052–4463. doi: 10.1038/s41597-021-00910-1.

[18] Moritz Buck. mOTUlizer: Github.com/moritzbuck/mOTUlizer, June 2021.

[19] Pierre-Alain Chaumeil, Aaron J Mussig, Philip Hugenholtz, and Donovan H Parks. GTDB-Tk: A toolkit to classify genomes with the Genome Taxonomy Database. Bioinformatics, 36(6):1925–1927, March 2020. ISSN 1367–4803. doi: 10.1093/bioinformatics/btz848.

[20] Maria G. Pachiadaki, Julia M. Brown, Joseph Brown, Oliver Bezuidt, Paul M. Berube, Steven J. Biller, Nicole J. Poulton, Michael D. Burkart, James J. La Clair, Sallie W. Chisholm, and Ramunas Stepanauskas. Charting the Complexity of the Marine Microbiome through Single-Cell Genomics. Cell, 179(7):1623–1635.e11, December 2019. ISSN 0092-8674, 1097-4172. doi: 10.1016/j.cell.2019.11.017.

[21] Martin Steinegger and Johannes Söding. MMseqs2 enables sensitive protein sequence searching for the analysis of massive data sets. Nature Biotechnology, 35(11):1026–1028, November 2017. ISSN 1546–1696. doi: 10.1038/nbt.3988.

[22] Carlos P. Cantalapiedra, Ana Hernández-Plaza, Ivica Letunic, Peer Bork, and Jaime Huerta-Cepas. eggNOG-mapper v2: Functional Annotation, Orthology Assignments, and Domain Prediction at the Metagenomic Scale. bioRxiv, page 2021.06.03.446934, June 2021. doi: 10.1101/2021.06.03.446934.

